# Neurophysiological Markers of Cancer-Related Fatigue Derived from High-Density EEG

**DOI:** 10.1101/2025.04.29.651322

**Authors:** Vikram Shenoy Handiru, Easter S. Suviseshamuthu, Haiyan Su, Guang H Yue

## Abstract

Cancer-related fatigue (CRF) significantly diminishes the quality of life of cancer survivors; however, objective diagnostic markers and the underlying neurophysiological mechanisms remain unclear. This study aimed to identify noninvasive EEG-based biomarkers of CRF by examining cortical activity and functional connectivity.

We recorded resting-state and task-related [repetitive submaximal elbow flexions (EFs) until self-perceived exhaustion] high-density electroencephalography (EEG) from 10 cancer survivors with CRF and 14 healthy controls (HC). In our analysis, task-induced fatigue was categorized as mild, moderate, and severe, corresponding to the level of fatigue perceived at the beginning, middle, and end of the task period.

Our study revealed the following significant findings: (1) Linear mixed-effects modeling of event-related desynchronization (ERD) EEG analysis during the EF task demonstrated significant effects of group and fatigue levels in the alpha band (8-12 Hz). (2) EF task-specific functional connectivity was estimated using the debiased weighted phase-lag index (dwPLI), which demonstrated reduced inter-regional connectivity in the M1 and prefrontal regions in the CRF group compared with the HC group. (3) The dwPLI analysis identified significantly reduced alpha-band connectivity strength in the CRF group, particularly between the right supramarginal gyrus and other brain regions during mild fatigue. (4) Additionally, resting-state EEG exhibited globally elevated delta-band (1-4 Hz) activity in CRF survivors than HC, potentially reflecting chronic fatigue. These observations emphasize the clinical relevance of resting-state EEG, motor activity-related ERD and functional brain connectivity as potential CRF biomarkers. Future research should validate these findings in larger cohorts and provide insights into more objective CRF diagnosis and the development of personalized interventions for alleviating CRF.

## 1 Introduction

Cancer-related fatigue (CRF) is one of the most distressing and prevalent symptoms experienced by cancer patients, affecting more than 70% of individuals undergoing treatment and often persisting long after its completion [1, 2]. Unlike ordinary fatigue, CRF is disproportionate to physical activity and is not relieved by rest, which significantly disrupts physical, emotional, and cognitive functioning. This debilitating condition affects the quality of life, impacting patients’ ability to work, maintain social relationships, and engage in daily activities [3, 4]. Despite decades of research, the multi-factorial nature of CRF—encompassing biological, psychological, and social dimensions—continues to pose significant hurdles for effective diagnosis, management, and treatment. Moreover, it is crucial to delineate the subtypes of fatigue, including but not limited to CRF-induced physical and mental fatigue. To this end, a review article by De Raaf et al. [2] emphasized the need for further research on the pathophysiological mechanisms underlying physical and mental fatigue. They noted that these two subtypes may represent separate symptoms associated with distinct neural markers and treatment responses. Therefore, this study focused on physical task-induced fatigue to identify neural biomarkers (while also examining global brain activity in a resting state) by analyzing high-density electroencephalography (EEG) data in cancer survivors exhibiting CRF.

Both the central and peripheral nervous systems play pivotal roles in physical fatigue, which is a key component of CRF [5–7]. In a review article by Bower et al., [1], inflammation was identified as a key mechanism underlying fatigue, wherein elevated proinflammatory cytokines due to cancer can signal to the central nervous system (CNS), leading to symptoms of fatigue. Central fatigue, governed by the CNS, alters cortical excitability, connectivity, descending command, and neurochemical signaling. In contrast, peripheral fatigue is characterized by reduced muscular force generation, which is often linked to impaired neuromuscular transmission and inefficient metabolism [8]. High-density electroencephalography (EEG) and electromyography (EMG) studies have provided critical insights into these processes. For instance, a study utilizing EEG and EMG recordings during a submaximal isometric elbow contraction task in cancer survivors reported a prominent role of central fatigue, evidenced by reduced voluntary muscle recruitment measured using evoked twitch force and a shorter endurance time compared to matched healthy controls (HC) despite lower levels of muscle fatigue [9]. Other EEG studies have also reported altered cortical activity during prolonged motor tasks, particularly in motor-related areas. EMG studies complement these findings by elucidating muscle activation patterns and fatigue-related declines in force output [10].

Zinn et al. evaluated the EEG source-space functional connectivity (FC) of the cortical autonomic network (CAN) in myalgic encephalomyelitis/chronic fatigue syndrome (CFS) to investigate alterations in the functional networks of physical fatigue due to pathological conditions. They noticed the following significant disruptions in both local and global network integration, measured by the graph-theoretic metrics of FC [11]. Connectivity in brain regions associated with somatomotor, cognitive, and affective symptom clusters is reduced, which emphasizes the role of CAN dysfunction in manifesting fatigue and related symptoms. Several other resting-state EEG studies have reported hyperactivity or higher delta-, theta-, and alpha-band power in the frontal and prefrontal regions in individuals with pathological fatigue conditions such as CFS [11, 12] and fibromyalgia syndrome [13] than in HC. These studies [5, 11– 13] have led to a current understanding of the brain mechanisms of *trait fatigue* in pathological conditions, which is persistent and independent of motor exertion. However, it is crucial to characterize *state fatigue*, which varies with the voluntary effort exerted during an activity (e.g., submaximal elbow contraction). Furthermore, oscillatory rhythms of brain activity play an essential role in characterizing task-relevant cortical activity and functional connectivity. To this end, we postulate that phase-based FC metrics and event-related desynchronization (ERD) measured for different EEG frequency bands will provide more insights into the neural mechanisms of CRF. Significant knowledge gaps remain despite our current understanding of altered brain activity and connectivity due to *state* and *trait* fatigue. For example, fMRI studies have demonstrated altered connectivity between the prefrontal cortex and motor regions, implicating these networks in fatigue-related motor and cognitive dysfunctions. However, the temporal resolution of fMRI limits its ability to capture real-time brain dynamics during fatigue-inducing tasks, and its relatively high cost hinders its widespread use. Therefore, we conducted a high-density EEG source-space analysis to study altered cortical activity and FC during rest and a submaximal isometric elbow flexion task until perceived exhaustion. Thus, we aimed to investigate the brain dynamics underlying CRF by computing EEG source-space cortical activity pertaining to the resting state and progressive fatigue levels at various time instants. Moreover, we studied task-relevant FC changes between the groups. Specifically, we aimed to assess two potential neural biomarkers of CRF: (1) EEG band power features, reflecting fatigue-induced changes in cortical activity and (2) FC estimated using the debiased weighted phase-lag index (dwPLI), reflecting alterations in brain network communication during task performance.

## 2 Methods

### 2.1 Participants

Participants included individuals diagnosed with CRF and matched HC. Patients with cancer were eligible if they had been off chemotherapy and radiation for at least four weeks, had a hemoglobin level greater than 10 g/dL, did not exhibit clinical depression or received antidepressants. Exclusion criteria included unintentional weight loss, severe pulmonary compromise requiring oxygen, or neuromuscular conditions affecting motor function. HC were matched by age, gender, and body mass index (BMI) and underwent similar screening procedures to exclude depression. Further details on participant characteristics and eligibility criteria can be found in [9]. In this study, the EEG data from 24 participants [HC (*n* = 14) and individuals with CRF (*n* = 10)] were analyzed.

### 2.2 Experimental Task

Participants performed submaximal intermittent contractions with elbow flexor muscles until they experienced severe fatigue. Each intermittent contraction was set at 40% of the individual’s maximal voluntary contraction (MVC) for a period of 5 s succeeded by a rest period of 2 s. The 40% MVC level was chosen to induce severe fatigue within 30 min according to the findings of Cai et al. [14]. Visual cues were provided on an oscilloscope screen at the start and end of each contraction. This task was repeated until participants reached self-perceived exhaustion.

### 2.3 EEG Data Recording and Preprocessing

EEG data were recorded using a high-density 128-channel system (Electrical Geodesics Inc., Eugene, OR, USA). The data were preprocessed and analyzed using EEGLAB (version 2021.0) and custom MATLAB scripts.

Raw EEG data were band-pass filtered with a passband of 1 Hz–50 Hz to remove noise. The artifact subspace reconstruction (ASR) procedure was then applied with the following parameter settings: (1) The rejection threshold for channel correlation was set to 0.8 to discard poorly correlated channels. (2) Channels with line noise standard deviation (SD) more than four times that of all channels were removed. (3) The ASR burst criterion was set to *k* = 20*×* to reconstruct epochs with abnormally high power, i.e., more than 20⨯SD of calibration data [15]. (4) The window criterion was set to 0.25 to remove excessively contaminated time windows—; if more than 25% of the channels were contaminated in a given time window, it was discarded. (5) The “bad” channels identified by ASR were corrected using spherical interpolation. The cleaned EEG data were decomposed using an independent component analysis (ICA) algorithm, namely, CUDA-ICA. The resulting independent components (ICs) were classified with the ICLabel technique; ICs representing brain activity with more than 75% probability were retained for further analysis. The data were re-referenced to the average reference.

Retained ICs were fitted using the standard boundary element model with the Montreal Neurological Institute (MNI) coordinate system. DIPFIT settings were applied, and dipoles were fitted for each IC. Outlier dipoles were removed based on a residual variance threshold of 20%, as implemented in our previous study [16] to ensure reliable dipole fitting.

Continuous EEG data were epoched from *−*1.5 s to 5 s around the auditory cue, marking the start of an intermittent elbow contraction trial. Epochs were designated to pertain to three fatigue levels (mild, moderate, and severe) based on their relative position within each session. Specifically, the first, middle, and the last 10% of the total number of epochs were deemed as the data associated with mild, moderate, and severe fatigue, respectively. Hereafter, mild, moderate, and severe fatigue are referred to as *MildFatg, ModFatg*, and *SevFatg*, respectively.

### 2.4 Brain Activity Analysis

To analyze the brain activity during graded fatigue levels, we computed the ERD power at various EEG frequency bands^1^ from EEG signals recorded from the electrodes overlying cortical regions associated with motor tasks. In this context, a few definitions related to the power spectral density analysis are provided below.

For simplicity, the subject index *n* and fatigue condition are omitted in (1)–(4). The mean event-related spectrum (ERSP) is the average data power spectrum across *M* trials computed for the sliding time windows centered at *t* in each trial, given by

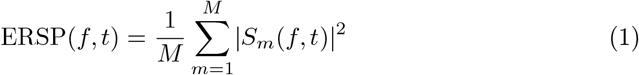

where *S*_*m*_(*f, t*) is the spectral estimate for trial *m* at frequency *f* and time instant *t*. The term |*S*_*m*_(*f, t*) | ^2^ represents the power spectrum for the *m*-th trial of subject *n*. For a given frequency, the absolute ERD was computed by dividing the ERSP power in (1) for the steady contraction at each time-frequency point by the average spectral power in the pre-contraction baseline period at the same frequency. ERD analyses are based on the log-transformed ERD in (2) and its time-averaged quantity in (3), both expressed in decibels (dBs):

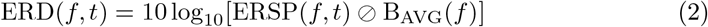

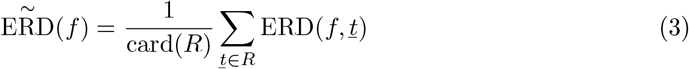

where

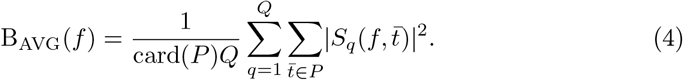

In (2)-(4), *⊘* represents Hadamard division; card(*R*) denotes the cardinality of the set of averaged steady-contraction time points in a fatigue condition; and B_AVG_(*f*) denotes the pre-contraction spectral power as a function of *f* averaged over the baseline-period time points *P* across *Q* trials, with *Q* being the total number of trials performed by a subject under all fatigue conditions. Thus, we derived a common baseline for every subject, i.e., the power spectra of the pre-contraction data for each subject were averaged across the trials, time points, and fatigue conditions. We vertically concatenated row vectors 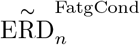 for *n* = 1, 2, …, *N* in each group (HC or CP) to construct a matrix of size *N × f* as given by

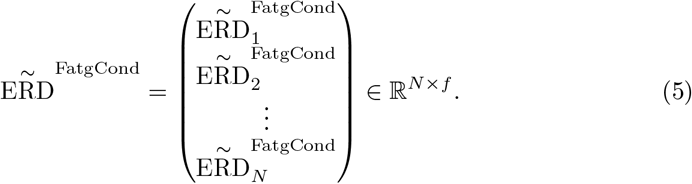

The power spectral density values in 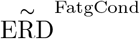 were averaged over the EEG frequency bands of interest.

### 2.5 EEG Source Imaging and Functional Connectivity Analysis

To circumvent the volume conduction effect encountered in scalp EEG analyses, we relied on EEG source estimates returned by a source-localization approach instead of EEG data recorded by the scalp electrodes. Note that EEG source estimation approaches are premised on the assumption that the scalp EEG signals are linear mixtures of the underlying cortical sources. Thus, the estimation of FC between different cortical regions of interest (ROIs) avoids spurious connections between EEG channels. We relied on MATLAB functions from the “ROI Connect” plugin developed for the EEGLAB toolbox [18] to compute a four-layered forward head model using the boundary element method [19] that models the brain, skull, and scalp surfaces with isotropic conductivity values. Since subject-specific MRIs were unavailable, we used template anatomy (ICBM152) based on a non-linear average of 152 adult MRI scans [20]. The functional connectivity between cortical ROIs was computed using a phase synchronization measure based on the weighted phase lag index (wPLI). Because the original form of the wPLI has a positive bias, we adopted the debiased wPLI proposed by

Vinck et al. [21], where the limitation was addressed.

**Primer**. The EEG data recorded from *L* number of EEG sensors or from *L* cortical source estimates per parcellated Desikan-Killiany atlas with *M* epochs (trials), each having *T* time points, is denoted as **E***∈* ℝ^*T ×L×M*^. By concatenating two arbitrary EEG channel recordings (or source estimates) of all the trials, the time-series data can be represented as a function of time, i.e., *x*_1_(*t*) and *x*_2_(*t*), *t* = 1, 2, …, *T × M*. The debiased wPLI between *x*_1_(*t*) and *x*_2_(*t*) can then be computed using their Fourier transforms, *X*_1_(*f*) and *X*_2_(*f*), respectively. By expressing the cross-spectrum between *X*_1_(*f*) and 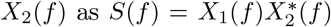,where indicates the complex conjugate, the debiased wPLI estimator (dwPLI) is given by

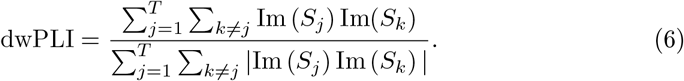

In (6), *S*_*j*_ and *S*_*k*_ denote the cross-spectrum between a pair of sources/sensors for the *k*-th and *j*-th time windows (or epochs), respectively; Im(*S*) represents the imaginary part of the cross-spectrum. Similar to other phase-based connectivity measures, dwPLI is robust in addressing the volume conduction effect.

We estimated the dwPLI for each EEG frequency band of interest, i.e., theta: = 4 Hz–8 Hz, alpha: = 8 Hz–13 Hz, and beta: = 13 Hz–30 Hz.

### 2.6 Statistical Analysis

To examine the effects of groups [HC (*N*_HC_ = 14) and CRF (*N*_CRF_ = 10)] and fatigue levels (mild, moderate, and severe) on ERD in the theta, alpha, and beta frequency bands, we conducted a linear mixed-effects (LME) model analysis in R. For ERD analyses, we averaged the time-frequency patterns from seven channels (FC3, FC1, C1, C3, C5, CP1, and CP3) overlying the contralateral motor region since we observed strong ERD activity patterns in alpha and beta band across participants as illustrated in Fig. 1. After formatting the ERD data using the ‘*tidyr* ‘package in R, separate LME models were fit for the alpha-and beta-band ERD using the ‘*lme4* ‘package. The models included fixed effects for the *Group* and *FatigueLevel* as well as a random intercept for each participant to capture individual variability. The model equations were specified as follows: alpha_ERD_ *∼ Group* + *FatigueLevel* + (1 |*participant*) and beta_ERD_ *∼ Group* + *FatigueLevel* + (1| *participant*). To analyze pairwise comparisons across fatigue levels for a given group, posthoc analyses were performed using estimated marginal means with the ‘*emmeans*’ package in R. Model summaries, including effect sizes and statistical significance, were reported using the ‘*lmerTest* ‘package in R.

**Fig. 1.**
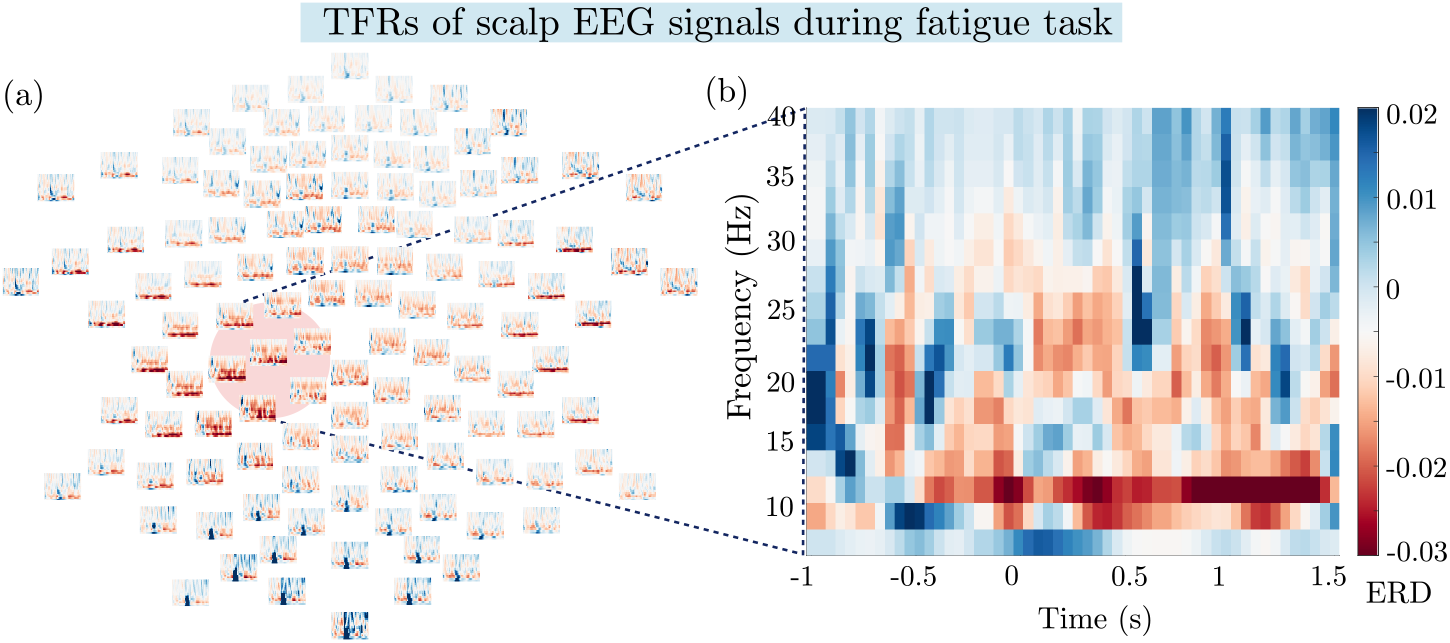
(a) Group-averaged time-frequency representations (TFRs) of EEG signals from all the scalp sensors during the fatigue task. (b) Averaged TFR corresponding to the EEG channels overlying the left sensorimotor region is magnified for better visualization.

To analyze the FC between 68 brain region parcellations per the Desikan-Killiany atlas, the dwPLI matrices of size 68 *×*68*× N*_Group_ were computed, where *N*_Group_ = 14 or 10, depending on whether the group under consideration is HC or CRF. The between-group comparisons (HC vs. CRF) were performed using a paired permutation-based multivariate *t*-test with multiple comparison corrections using the ‘*statcond* ‘function in EEGLAB. Statistical maps (*t*-maps and *p*-values) were computed for each frequency band and fatigue condition. Type I errors were controlled with the false discovery rate (FDR), and the significant connections were shown in the thresholded *t*-maps. To view the connectivity results in an anatomical context, cortical models were overlaid using the *BrainNet Viewer* (Xia et al., 2013). Group-level differences in dwPLI were mapped onto cortical surfaces with statistically significant FC values as edges and ROIs as nodes. Note that the *p*-values in the map were log-transformed to enhance the visualization.

## 3 Results

### 3.1 Demographic Characteristics

The demographic and baseline characteristics of the study participants are summarized in Table 1. The mean age of the participants was 53 *±* 11.2 years for the CRF group and 59.7 *±* 9.7 years for the HC group. Sex distribution was comparable between groups, with 40% males and 60% females in the CRF group and 42.86% males and 57.14% females in the HC group. The cancer staging varied within the CRF group, with a higher prevalence of advanced-stage cancer. Functional status, as measured by the Eastern Cooperative Oncology Group (ECOG) performance status, indicated an inability to engage in strenuous activity in the CRF group (mean ECOG score = 1.3 *±* 1.06). The endurance level (time until perceived exhaustion) and self-reported fatigue severity in resting state quantified by the Brief Fatigue Inventory (BFI), were significantly different between the groups, with the CRF group exhibiting lower endurance in performing the motor task (mean endurance = 342.3 *±* 135.1 s) and higher fatigue scores (mean BFI score = 4.5) relative to the HC group, whose mean endurance score was 531.7 *±* 137.2 s and the mean BFI score was 0.9 *±* 1 (Table 1).

**Table 1.** Summary Statistics of the Study Demographics.

| Characteristic | Total ($n = 24$) | HC ($n = 14$) | CRF ($n = 10$) | Group difference ($p$-values) |
| --- | --- | --- | --- | --- |
| Age (years) | $55.9 \pm 10.9$ | $53.0 \pm 11.2$ | $59.7 \pm 9.7$ | 0.14 |
| BFI | $2.5 \pm 2.3$ | $0.9 \pm 1.0$ | $4.5 \pm 1.9$ | $1.6e - 05$ |
| Endurance (s) | $445.6 \pm 164.3$ | $531.7 \pm 137.2$ | $342.3 \pm 135.1$ | 0.004 |

### 3.2 Group Comparison of ERD Features Using LME Models

We conducted an analysis with the LME model using restricted maximum likelihood estimation to investigate the effects of the *Group* and *FatigueLevel* on the ERD as the dependent variable. The model included a random effect for subjects and was standardized for parameter estimation. Confidence intervals (95% CIs) and *p*-values were derived using a Wald *t*-distribution approximation. The model’s total explanatory power was substantial, with a conditional *R*^2^ of 0.59 and a marginal *R*^2^ of 0.30 for the fixed effects alone.

For alpha ERD, the model revealed a significant main effect of *Group*, indicating that the ERD of the HC group was significantly higher (less negative) than that of the CRF group (*β* = 1.151, SE = 0.527, *t* = 2.184, *p* = 0.04). There was also a significant effect of *Fatigue Level* —with both ModFatg (*β* = 1.507, *p <* 0.001) and SevFatg (*β* = 2.418, *p <* 0.001) showing higher ERD compared to MildFatg. By controlling for *Group* and applying Tukey’s adjustment, each pairwise comparison among the three fatigue levels showed a significant difference in alpha ERD. MildFatg had a significantly lower alpha ERD than ModFatg (*t* = *−*4.219, *p* = 0.0003) and SevFatg (*t* = *−*6.772, *p <* 0.0001). Additionally, ModFatg had a significantly lower alpha ERD than SevFatg (*t* = *−*2.553, *p* = 0.0368). The estimated residual variance was 1.530 (SE = 1.237). The intra-class correlation was 0.421, suggesting that individual differences contribute meaningfully to the overall variation in ERD values.

Similarly, a marginally significant main effect of *Group* was observed for beta ERD, suggesting a trend toward higher ERD values in the HC than the CRF group (*β* = 0.789, SE = 0.417, *t* = 1.894, *p* = 0.07). A significant effect of *Fatigue Level* was found, i.e., both ModFatg (*β* = 0.999, *p <* 0.001) and SevFatg (*β* = 1.780, *p <* 0.001) have a significantly higher beta ERD than MildFatg. By controlling for *Group* and applying Tukey’s adjustment, each pairwise comparison among the three fatigue levels revealed a significant difference in beta ERD as follows. MildFatg had a significantly lower beta ERD than ModFatg (*t* = *−*3.583, *p* = 0.0023) and SevFatg (*t* = –6.380, *p <* 0.0001). Furthermore, ModFatg had a significantly lower beta ERD than SevFatg (*t* = *−*2.797, *p* = 0.0202). The estimated residual variance was 0.934 (SE = 0.966), and the intra-class correlation was 0.429, suggesting individual differences contribute meaningfully to the variance of ERD values. The results from both the alpha-and beta-band LME models are concisely presented in Table 2 and summarized in Fig. 2. As summarized in Table 2, both the main effects, i.e., *Group* and *FatigueLevel*, were significantly different in alpha-band ERD. However, the effect of *Group* was not statistically significant (*p* = 0.07) for the beta-band ERD.

**Table 2.**
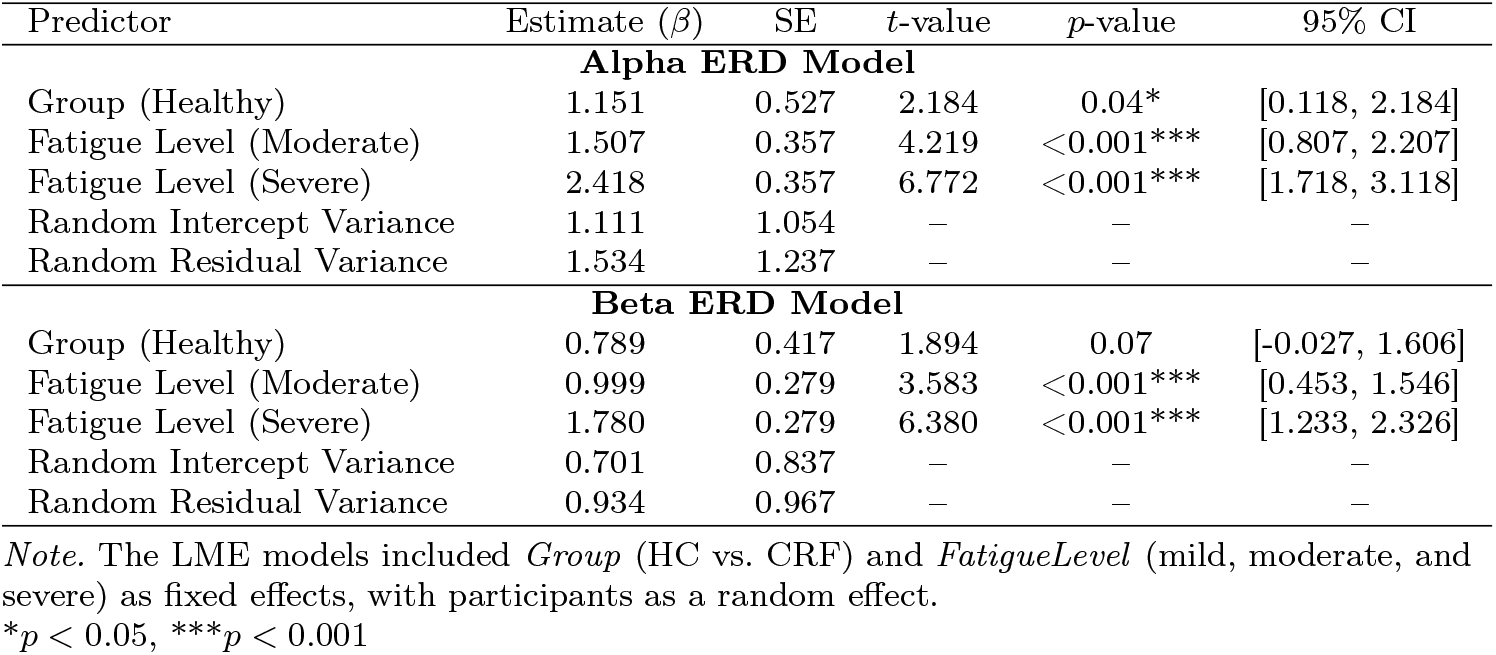
LME Model Results for Alpha and Beta ERD.

**Fig. 2.**
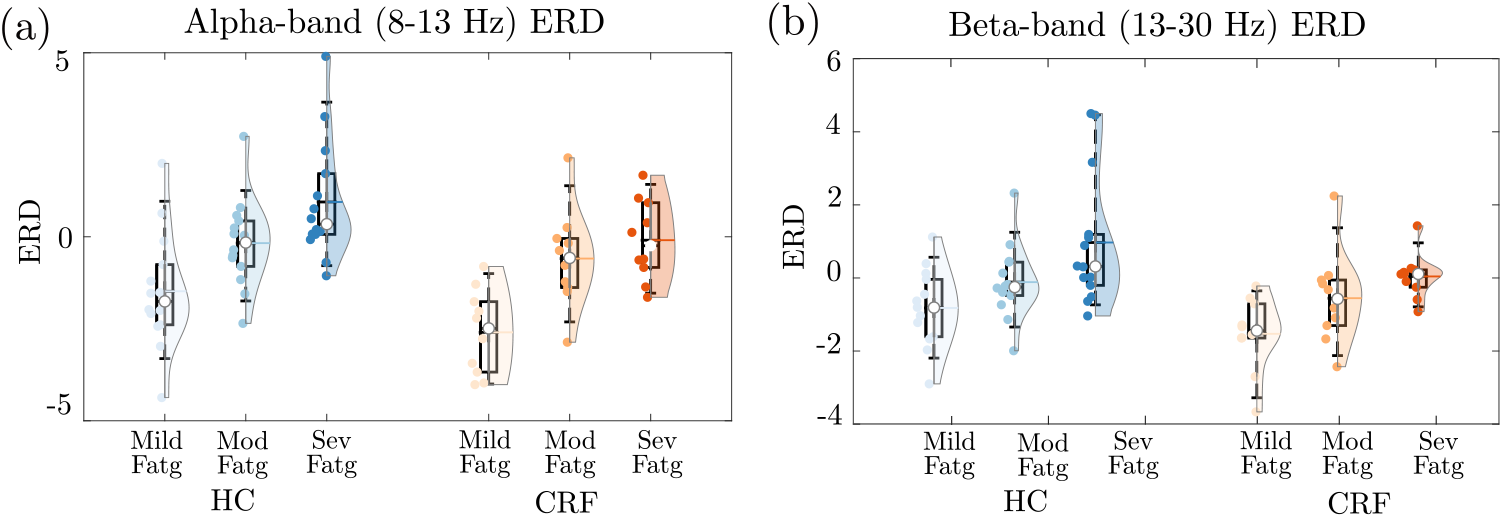
Half violin plots of (a) alpha– and (b) beta-band ERD values for left M1 across different fatigue levels (mild, moderate, and severe) and groups (HC in blue and CRF in orange). The shaded regions represent the 95% confidence intervals.

#### Resting-state EEG

Results for the cortical network-level spectral power differences are displayed using bar plots within each group for the resting-state EEG in the delta band (1-4 Hz) in Fig. 3. Notably, the delta-band power was higher in the CRF group than in the HC group across all the cortical networks. There were no noticeable differences within the group across the networks.

**Fig. 3.**
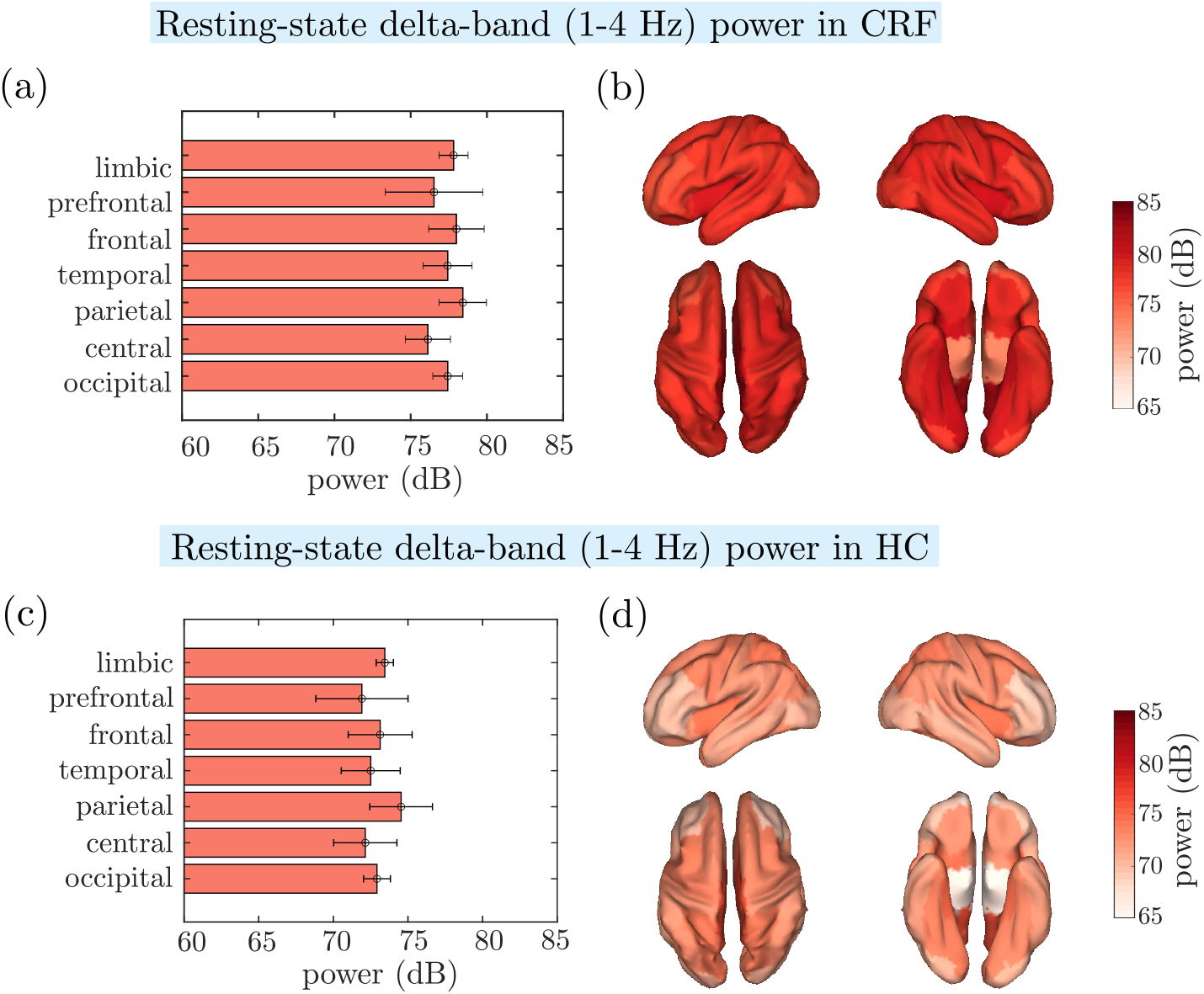
Horizontal bar plots of the resting-state delta-band power of cortical regions grouped by functional networks in the (a) CRF and (c) HC group. Delta-band power mapped to the cortical sources estimated with the linear constrained minimum-variance beamformer is displayed for (b) CRF and (d) HC group. In (b) and (d), the left-right lateral and dorsal-ventral views of the cortical surface are presented in the top and bottom rows, respectively.

**Fig. 4.**
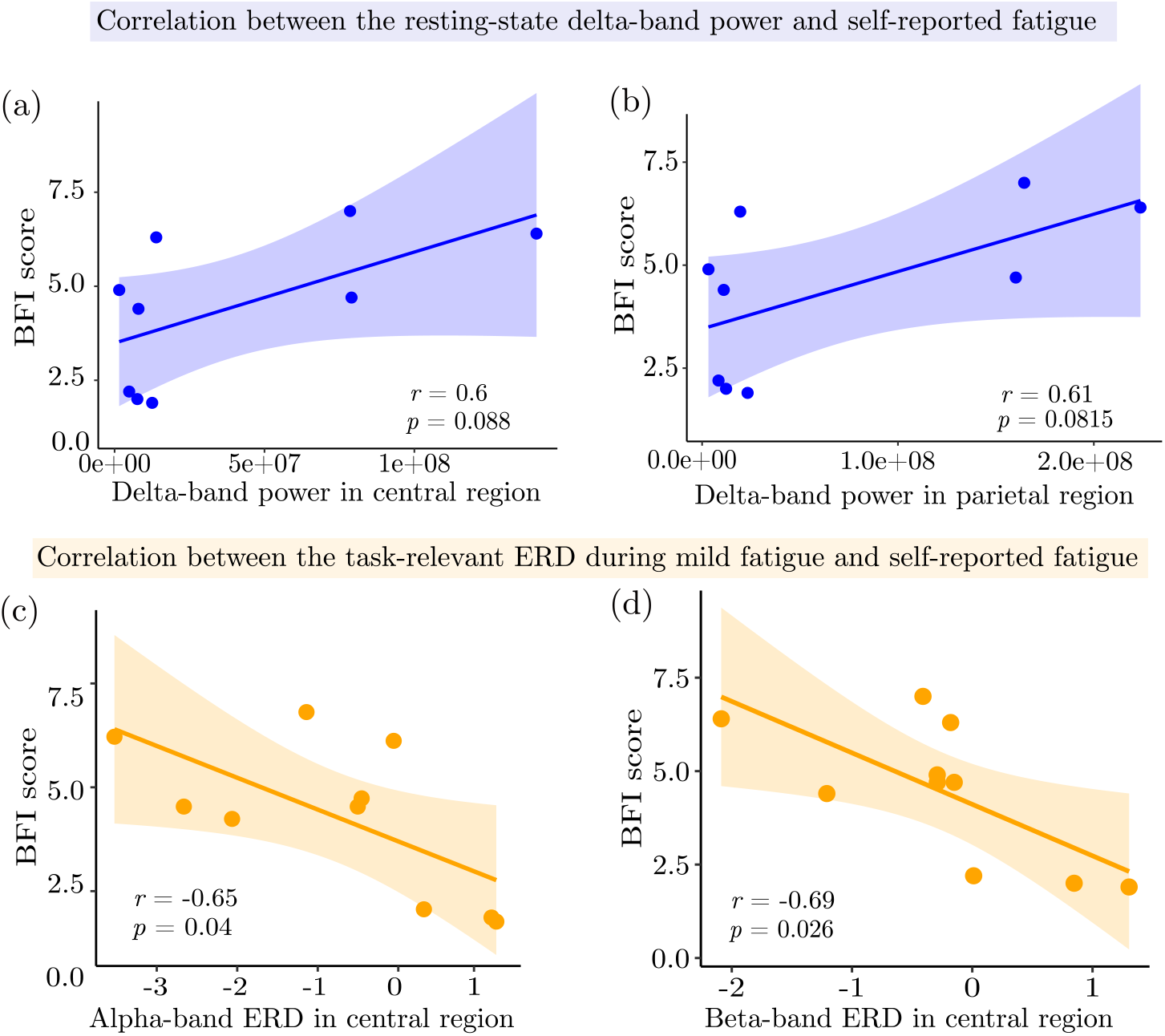
Scatter plots that show the correlation between the following band-specific power values and BFI in the CRF group: resting-state delta-band power of the (a) central and (b) parietal region; task-related (MildFatg) (c) alpha and (d) beta ERD of the central region. Each point represents a participant, with solid lines and shaded areas indicating the best-fit linear regressions and 95% confidence intervals, respectively. The Pearson correlation coefficient (*r*) and *p*-value are displayed within each panel.

We performed a correlation analysis to observe the relationship between the self-reported fatigue measure at rest (BFI score) and the delta-band power from the resting-state EEG. For comparison, we also reported a correlation between the BFI score and alpha-/beta-band ERD during MildFatg. At rest, the delta-band power of the central and parietal regions was positively correlated with the BFI score, although it was not statistically significant. Pearson’s product-moment correlation between the delta-band power of the central region (sensorimotor areas) and BFI was *r* = 0.6, *p* = 0.088, while that between the delta-band power of parietal areas and BFI was *r* = 0.61, *p* = 0.0815. In contrast, during MildFatg, the alpha-and beta-band ERD values of the central region were negatively correlated with BFI [22], i.e., *r* = *−*0.65, 95% CI [*−*0.91, *−*0.04], *t* = *−*2.44, *p* = 0.040 and *r* = *−*0.69, 95% CI [*−*0.92, *−*0.11], *t* = *−*2.73, *p* = 0.026, respectively.

### 3.3 Group Comparison of Functional Connectivity Patterns

Multivariate *t*-test analysis yielded a *t*-map and a *p*-map, both of size 68*×* 68. The results are presented in Figures 5 and 6, respectively. In these figures, the *t*-map provides the *t*-statistic values for the connectivity differences between groups. Positive values (in blue) indicate higher connectivity in the HC group, whereas negative values (in red) indicate higher connectivity in the CRF group. Likewise, the *p*-map provides the *p*-values corresponding to the *t*-statistics, indicating the statistical significance of the observed differences. Given the multivariate comparison, only the *p*-values below a predefined threshold (e.g., *p <* 0.05) after false discovery rate (FDR) correction were considered statistically significant. To visualize the *p*-map, we have log-transformed the p-values as *−*log_10_(*p*), so that *p* = 0.05 and 0.01 are represented as 1.301 and 2, respectively.

**Fig. 5.**
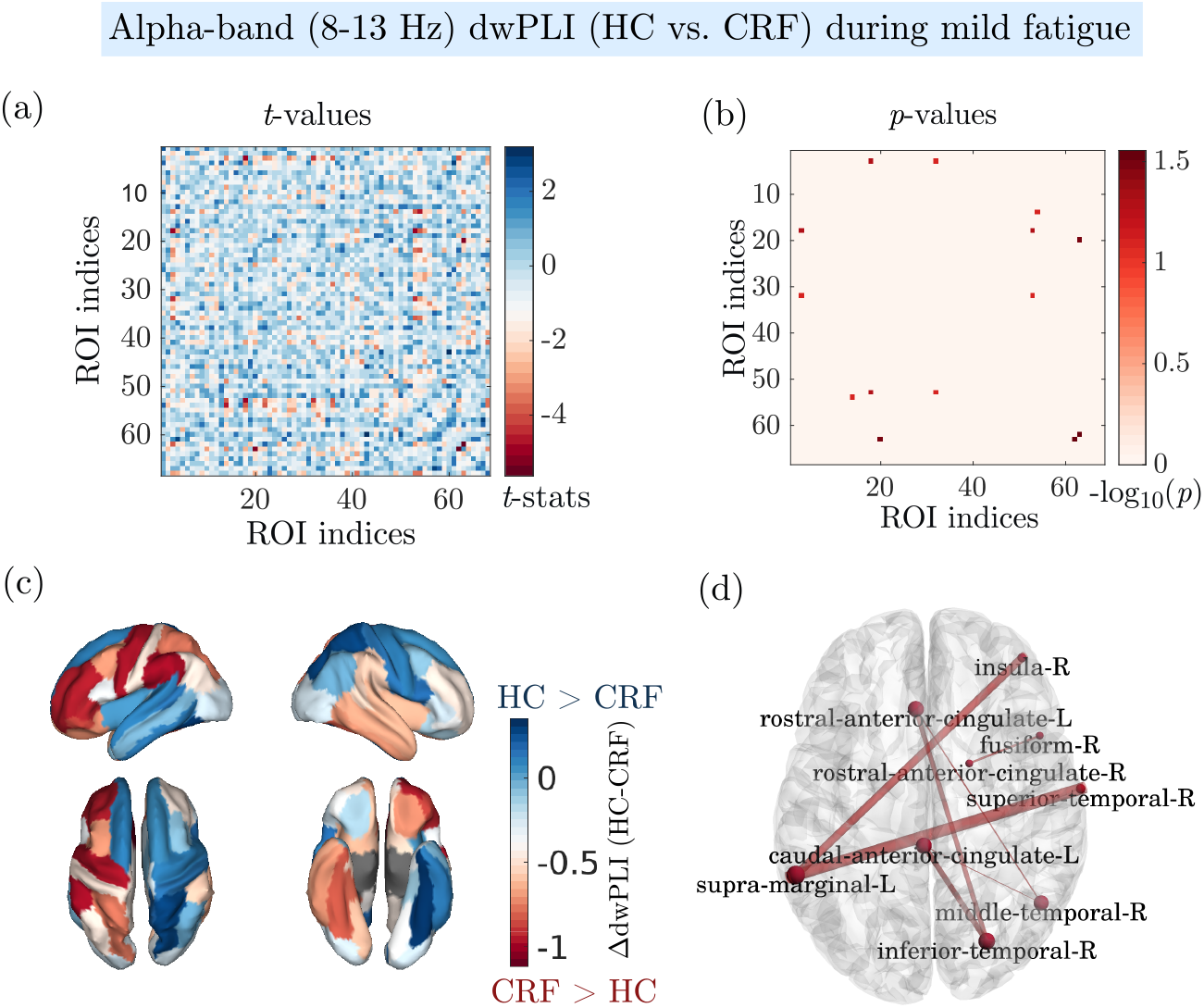
Alpha-band functional connectivity based on dwPLI for HC and CRF groups during mild fatigue. (a) Heatmap displaying *t*-values for pairwise comparisons of the dwPLI between ROIs. Here, the ROI indices on the x and y axes correspond to the Desikan-Killiany atlas-based 68 anatomical parcels sorted alphabetically. (b) Corresponding *p*-values highlighting significant connections. (c) Pictorial representation of *t*-statistics-based differences in dwPLI strength between HC and CRF across the cortical surface. Blue regions indicate higher dwPLI in HC than CRF, while red regions imply the opposite. (d) Brain network topology shows significantly different connections between ROIs in the two groups, with CRF exhibiting greater connectivity for all ROI pairs than the HC group. The thickness of the lines represents the magnitude of the difference in connectivity strength.

**Fig. 6.**
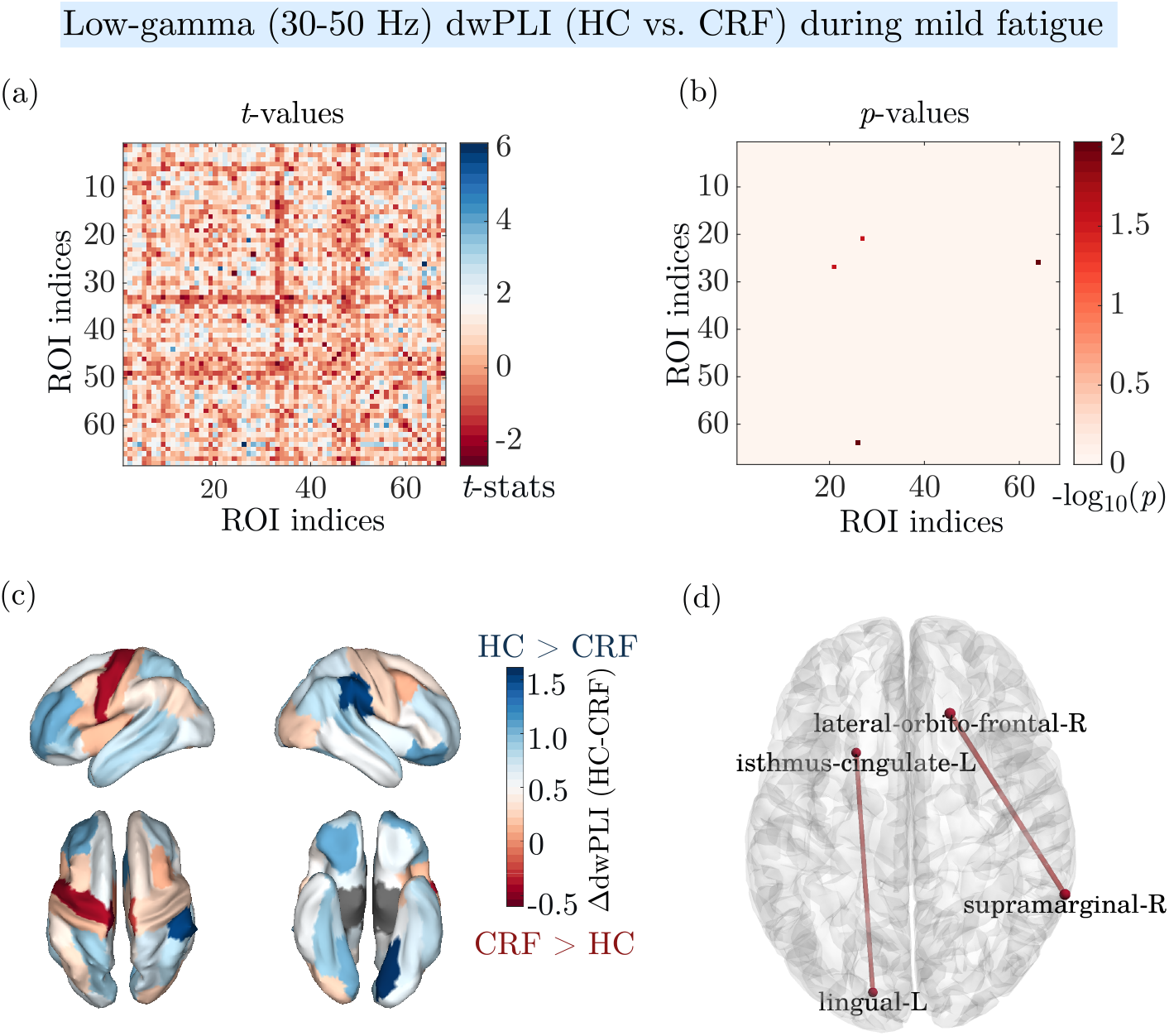
Low-gamma band functional connectivity based on dwPLI for HC and CRF groups during mild fatigue. (a) Heatmap displaying *t*-values for pairwise comparisons of the dwPLI between ROIs. (b) Corresponding *p*-values highlighting significant connections. (c) Pictorial representation of *t*-statistics-based differences in dwPLI strength between HC and CRF across the cortical surface. Blue regions indicate higher dwPLI in HC than CRF, while red regions imply the opposite. (d) Brain network topology depicts significantly different connections between ROIs in two groups, with CRF exhibiting greater connectivity for the two ROI pairs than the HC group. The thickness of the lines represents the magnitude of the difference in connectivity strength.

The unpaired *t*-test comparison of functional connectivity strengths in the alpha-band revealed significant between-group differences in several brain regions. The regions with uncorrected *p*-values less than 0.05 included the following in the left hemisphere: caudal anterior cingulate cortex (*t* = *−*2.558, *p* = 0.018), caudal middle frontal gyrus (*t* = *−*2.233, *p* = 0.039), parstriangularis (*t* = *−*2.328, *p* = 0.034), precentral gyrus (*t* = *−*2.462, *p* = 0.022), rostral anterior cingulate cortex (*t* = *−*2.630, *p* = 0.016), rostral middle frontal gyrus (*t* = *−*2.353, *p* = 0.029), and supramarginal gyrus (*t* = *−*2.256, *p* = 0.039). However, it must be noted that these network strength differences did not survive the FDR correction. A similar analysis conducted for the low-gamma band (31-60 Hz) during MildFatg condition revealed that only the right supramarginal gyrus had a significantly higher connectivity strength in HC compared to CRF, after accounting for the FDR correction [*t* = 4.122, *p* = 0.00055 (uncorrected), *p* = 0.037 (FDR-corrected)].

## 4 Discussion

The findings from the fatigue EEG data point to significantly greater fatigue in both the resting state (BFI score) and after physical activity [sustaining elbow flexion (EF) contractions] in cancer survivors than in HC. These fatigue results are discussed in more detail in [9].

The primary goal of the current study was to investigate the neurophysiological variables in cancer survivors and HC in the resting state (band-specific EEG power) and during the EF fatiguing task (band-specific ERD and EEG-based FC between ROIs) for identifying potential neurophysiological markers of CRF. Our results revealed the following distinct EEG oscillatory and connectivity patterns associated with CRF: (1) The global delta-band EEG power during the resting state was notably higher in the CRF than in the HC group. (2) The self-perceived fatigue measure (BFI score) was positively correlated (not statistically significant) with the delta-band EEG power recorded from both the central and parietal regions in the CRF group during the resting state. (3) The BFI score and the alpha-or beta-band EEG power were negatively correlated during mild physical fatigue in the CRF group. (4) Several ROIs exhibited stronger functional connectivity during mild physical fatigue in the CRF but not in the HC group. In addition to these unique CRF-related neurophysiological findings, we observed that the alpha-and beta-band EEG power was scaling with the level of physical fatigue in both groups.

### 4.1 Delta-band EEG power in resting state

Our EEG power analysis of the resting state demonstrated significantly higher global (involving all ROIs) delta-band power in the CRF group than in the age-matched HC group (Fig. 3). To the best of our knowledge, this observation is unique. Given that CRF measurements in most studies are conducted at rest and cancer survivors with CRF report fatigue and tiredness that limit their ability to engage in physical-, recreational– and work-related activities, this finding provides a neurophysiological basis for the subjective feeling of fatigue. The hyperactive delta activity in a global fashion may suggest that this delta activity is so strong that it makes all other brain regions resonate with this frequency and perhaps generates a feeling of tiredness and fatigue even in the resting state. Thus, global delta-band power may reflect a feeling of fatigue in cancer survivors and potentially serve as a neurophysiological marker of CRF, especially at rest during everyday life.

### 4.2 EEG activity patterns during fatiguing task

In the frequency domain, we observed a significant increase in alpha-band power in the contralateral sensorimotor cortex, which is consistent with a previous study [23]. The authors reported that the alpha– and beta-band power increased in specific cortical regions during a fatiguing handgrip task, particularly in Brodmann area 40 (BA40), suggesting its role in integrating fatigue-related sensorimotor information. While we found an increase in alpha– and beta-band power in the precentral gyrus in agreement with [23], we also observed an increase in task-specific power in the postcentral gyrus, contrary to the findings of [23]. This discrepancy may be due to the study population, as our study examined CRF across multiple conditions (within-group and between-group comparisons), whereas the focus of [23] was healthy individuals in a crossover (fatigue vs. control) design.

### 4.3 Relationship trend between EEG spectral power and BFI

In addition to an increased global delta-band activity, the delta-band power in the central or parietal region enclosing sensorimotor cortices and BFI score showed a positive correlation trend [*p* = 0.088 and 0.082, respectively; Fig. 4 (a) and (b)] in the resting state of the CRF group. Park et al. (2019) [24] reported a significant positive correlation between frontal delta-band power and fatigue severity score in both CRF and chronic fatigue syndrome populations, reinforcing the correlation between slow-wave (delta) activity and fatigue perception. Similarly, Moore et al. [25] found that post-exertion EEG spectral power, including delta, increased in fatigued cancer patients during chemotherapy. These findings may be related to elevated delta wave activity due to disrupted hypothalamic-pituitary-adrenal axis, a mechanism observed in transdiagnostic fatigue [5].

The significant negative correlation between the alpha-or beta-band ERD and BFI [*p* = 0.04 and 0.026, respectively; Fig. 4 (c) & (d)] during mild physical fatigue may indicate a neurophysiological link between the baseline subjective fatigue state and subsequent task-induced changes in motor cortical activity [26]. In other words, we postulate that individuals experiencing higher resting-state fatigue enter physically demanding tasks with fewer cortical resources, potentially resulting in an increased perception of fatigue as the task progresses. Prior studies have shown a pre-fatigue-to-post-fatigue increase in alpha and beta oscillatory activities during elbow contractions, although these studies focused on healthy populations rather than individuals with CRF [27, 28]. These findings provide a context for interpreting the similar fatigue-related EEG changes observed in our CRF-focused study.

### 4.4 Functional connectivity between brain regions during fatiguing task

FC analysis during physical activity revealed notable alterations associated with CRF. Specifically, we obtained three key findings: (1) overall reduced inter-regional FC strength between the right supramarginal gyrus and other brain regions, (2) increased connectivity strength between the precentral gyrus and other cortical regions in CRF relative to HC in alpha and low-gamma frequency bands, and (3) no significant within-group differences across fatigue conditions after multiple-comparison correction.

The observed hyperconnectivity in CRF, particularly involving the precentral gyrus and prefrontal regions, suggests the compensatory recruitment of motor and cognitive control networks during physically demanding tasks. Previous studies [6, 29, 30] have consistently demonstrated the role of the precentral gyrus (primary motor cortex, M1) in voluntary motor output regulation and muscle force modulation during fatigue. The precentral gyrus facilitates sensorimotor integration, critical for generating motor commands and adjusting muscle performance as fatigue progresses [23]. Enhanced connectivity with prefrontal regions such as the anterior cingulate cortex (ACC) and rostral middle frontal gyrus in CRF likely indicates compensatory recruitment of these higher-order cognitive regions involved in effort perception, error monitoring, and motivation to sustain task performance despite fatigue [29, 31]. Jiang et al. [6] reported strengthened functional connectivity between the prefrontal and motor cortices in fatigue conditions, suggesting that such integration could facilitate sustained voluntary effort despite peripheral muscle fatigue.

The supramarginal gyrus (SMG), another region that shows significant connectivity changes, is involved in proprioceptive and somatosensory integration, which is crucial for maintaining precise motor control and detecting physical fatigue [23]. Increased alpha-band connectivity involving the SMG in CRF during mild fatigue may reflect impaired sensorimotor integration during the early stages of fatigue perception. However, we did not find any differential patterns of SMG in the severe fatigue stage, which should be studied in the future.

In line with findings from fMRI-based connectivity studies in CRF [32–34], altered FC in large-scale networks, including the default mode network (DMN), salience network, and dorsal attention network, was observed. For instance, the regions belonging to the DMN—involved in internal attention and self-referential processing [33, 34]—showed reduced coupling strength of low-gamma FC in the CRF group compared to HC, suggesting a compensatory mechanism to cope with fatigue. In contrast, impaired alpha-band FC in regions such as the ACC, a key node of the salience network, suggests maladaptive fatigue perception [35]. This maladaptive pattern could explain why cancer survivors perceive elevated fatigue levels even at rest.

### 4.5 Neurophysiological processes underlying physical fatigue in CRF

One notable observation in our study was the scaling of alpha-band ERD in the contralateral motor cortex with the level of fatigue. This trend aligns with those of previous studies [36, 37], which explains the involvement of M1 during physical fatiguing tasks. Specifically, Liu et al. [36] demonstrated that cortical activity in the sensorimotor cortex intensifies as motor unit recruitment increases to compensate for force loss due to muscle fatigue. Our findings support this notion, suggesting that individuals with CRF compensate by increasing motor cortical drive to sustain muscle output and task performance.

This compensatory mechanism, particularly during fatiguing motor tasks, is referred to as *supraspinal facilitation* [38]. As per the authors, as fatigue progresses, inhibitory inputs to M1 initially limit motor unit recruitment—a mechanism mediated by sensory feedback from fatigued muscles through small-diameter afferents (groups III and IV sensory nerves). However, voluntary effort can override this inhibition, enabling the motor cortex to increase the descending corticomuscular drive to sustain the task. This dynamic shift between early inhibitory responses and subsequent supraspinal facilitation is a hallmark of fatigue-related neural adaptations. Neuroimaging studies have demonstrated continued increases in cortical activity within the prefrontal cortex (PFC), cingulate gyrus (CG), and supplementary motor area (SMA), even as the activity in the contralateral M1 began to decline, suggesting a compensatory role for these regions in reinforcing cortical motor output during muscle fatigue [36]. This postulation is supported by the significantly strengthened functional connectivity between the PFC (a higher-order brain region) and contralateral M1 during severe muscle fatigue [31]. Such compensatory strategies are less pronounced in resting-state studies focusing on trait fatigue.

## 5 Limitations and Future Directions

Although our study provides novel insights into the EEG-based neural underpinnings of CRF, several limitations must be acknowledged. First, our sample size was relatively small, which may have limited the statistical power to detect finer connectivity differences. Second, the self-reported fatigue score measured using the BFI is prone to floor effects, particularly in the HC group, thus reducing the sensitivity in distinguishing fatigue-related variations. Third, our study did not involve subcortical structures such as the thalamus and basal ganglia, which may play a critical role in CRF pathophysiology.

Some of these issues can be addressed as follows: Future research should assess the reliability and validity of fatigue markers by conducting multi-session studies in larger cohorts. Since differences in connectivity patterns between the HC and CRF groups may partly be attributed to altered thalamocortical interactions, further investigation using multimodal imaging approaches [e.g., transcranial magnetic stimulation (TMS)– EEG or fMRI-EEG] may provide additional information.

Furthermore, we recommend that interventional studies focused on treating central fatigue incorporate multimodal imaging techniques to precisely target brain regions with disrupted cortical activity and FC. For example, noninvasive neuromodulation techniques such as transcranial direct current stimulation and repetitive TMS may be potential targeted intervention approaches. In support of this, a recent review highlighted the role of rTMS in reducing the pain and fatigue symptoms of chemotherapy-induced peripheral neuropathy—, a common side effect of cancer survivors undergoing chemotherapy [39]. On the premise of our valuable study findings and recent advances in neuromodulation, future research may explore EEG-guided targeted neuromodulation as an adjuvant to current pharmacological interventions. In conclusion, integrating these approaches could lead to a paradigm shift in CRF treatment, enhance fatigue symptom management, and improve overall quality of life in cancer survivors.

In conclusion, this study provides important insights into the neurophysiological mechanisms underlying CRF. We observed distinct resting-state EEG markers, notably increased global delta-band power in CRF participants compared to HC, potentially reflecting baseline fatigue perception. During motor fatigue tasks, individuals with CRF exhibited unique patterns of alpha– and beta-band ERD in sensorimotor areas and altered FC involving regions critical for sensorimotor integration and cognitive effort. Moreover, our correlation analysis revealed that higher self-reported fatigue was modestly associated with increased resting-state delta power and inversely related to task-induced alpha– and beta-band ERD, highlighting divergent neurophysiological patterns that may distinguish baseline fatigue perception from effort-induced cortical responses in CRF. These findings would enhance the understanding of the central mechanisms of CRF and suggest plausible EEG biomarkers to facilitate objective assessment and targeted therapeutic interventions in the future.

## Acknowledgements

This research was supported in part by grants from the National Institutes of Health (R01CA189665), Cleveland Clinic (RPC6700), and Department of Defense (DAMD17-01-1-0665) for data collection at Cleveland Clinic, and by the National Institute on Disability, Independent Living, and Rehabilitation Research (ARHF17000003) for data analysis conducted at Kessler Foundation. We also acknowledge the use of a generative language model—(ChatGPT OpenAI, GPT-4.5)—for manuscript refinement, language editing, and assistance with the abstract preparation.

## Ethics approval and consent to participate

This study involving human participants was reviewed and approved by the Institutional Review Board of the Cleveland Clinic. All participants provided written informed consent before their participation.

## Data availability statement

The datasets generated for this study are not publicly available due to institutional review board (IRB) restrictions protecting participant confidentiality. However, MATLAB scripts and de-identified preprocessed EEG data can be provided upon reasonable request and directed to the corresponding author, Guang Yue.

## Author contribution

VSH performed data analysis and drafted the initial manuscript. VSH and ESS jointly conducted the data analyses. GY conceptualized the study and secured research funding. HS provided guidance on the statistical analyses. All the authors contributed substantially to the review, revision, and approval of the final version of the manuscript.

## Footnotes

1 To obtain a physiologically meaningful interpretation of results, the channel-level ERD was computed for individual EEG frequency bands as each one has different functional characteristics. The typical frequency bands and their approximate spectral boundaries in adults [17] are as follows: delta – 1 Hz to 4 Hz; theta – 4 Hz to 8 Hz; alpha – 8 Hz to 13 Hz; beta – 13 Hz to 30 Hz; low gamma – 30 Hz to 50 Hz; and high gamma – 50 Hz to 80 Hz.

